# Refining Spatial Proteomics by Mass Spectrometry: An Efficient Workflow Tailored for Archival Tissue

**DOI:** 10.1101/2024.01.25.577263

**Authors:** Rune Daucke, Charlotte V. Rift, Nicolai S. Bager, Kartikey Saxena, Peter R. Koffeldt, Jakob Woessmann, Valdemaras Petrosius, Eric Santoni-Rugiu, Bjarne W. Kristensen, Pia Klausen, Erwin M. Schoof

## Abstract

**Background:** Formalin-fixed, paraffin-embedded (FFPE) tissue remains the gold standard for extensively archiving biological specimens, providing biobanks with large repositories of retrospective potential. However, while formalin crosslinking is effective at preserving tissue, it poses significant challenges for extracting molecular information, including the proteome. Traditionally, this process required high levels of input material, which, in turn, limited the ability to preserve cell-type heterogeneity and spatial information. To address these limitations, we developed an easily adaptable and highly efficient workflow for extracting deep proteomes from low-input materials, such as biopsies used in routine histopathological diagnostics.

**Methods:** We compared the extraction efficiency of pancreatic acinar cells identified in FFPE tissue samples stained with conventional hematoxylin-eosin (H&E) against that of cells isolated from tissue samples immunostained for the epithelial cell adhesion molecule (EpCAM) across material inputs ranging from 1,166 to 800,000 *µ*m^2^ (estimated to 2 to 1,310 cells in volume). Cells were isolated using laser capture microdissection and subsequently analyzed using Liquid Chromatography-Tandem Mass Spectrometry.

**Results:** Similar yields for both methods were observed, with EpCAM-positive cells yielding slightly higher results—approximately 1,200 unique protein groups at the lowest input and up to 5,900 at the highest. In cells isolated from H&E-stained tissue, ∼900 to ∼5,200 protein groups were identified. We decided that the optimal balance for our workflow, ensuring maximum protein identification while minimizing input material, lies within the range of approximately 50,000 to 100,000 *µ*m^2^. With these results, we tested spatial capabilities and biological relevance by isolating cancer cells from biopsies of pancreatic cancer, lung cancer, or glioblastoma, with the first two being stained with EpCAM and the latter being stained against the tumor-suppressor protein p53. We successfully identified tissue-specific protein expressions and observed prominent clustering of all cell populations.

**Discussion:** Our results highlight the feasibility of performing spatial proteomics on FFPE tissue using minimal input material. This adaptable methodology opens up possibilities for investigating cell-type-specific biology while preserving spatial and histological information.

## Introduction

With its ease of handling and cost-efficient storage options, formalin-fixed, paraffin-embedded (FFPE) remains the prevailing method for preserving biological specimens for use in histopathology, resulting in the establishment of extensive repositories at pathology departments worldwide. These repositories hold enormous untapped potential for conducting retrospective studies (1,2). Yet, a prevailing limiting factor for the use of FFPE tissue in proteomics has been the need for substantial sample quantities to acquire deep protein coverage, generally limiting proteomics to bulk analysis (3). This is highly problematic in the context of spatial proteomics, as it obscures the profiling of heterogeneous cell populations within tissues such as cancer (4-8). In addition, the limited availability of tissue, especially from small biopsies used in routine diagnostics or in rare diseases, can be too scarce to support comprehensive retrospective proteomic analyses.

The accelerated developments within the field of liquid chromatography-mass spectrometry (LC-MS) have recently enabled a level of sensitivity capable of analyzing ultra-low sample input (8,9), which consequently sparked a rapid emergence of methods transitioning from bulk or high-load input to low-load and even single-cell analysis (10-15). However, proteome coverage and ease of implementation on large-scale cohorts are still limited by sample preparation, which is especially true for FFPE tissue. To tackle these challenges, we developed an easily adaptable method for highly efficient protein extraction from laser capture microdissection-collected (LCM) FFPE samples. Our protocol builds on a similar approach introduced by Kwon et al., 2022 (10), where we collect the dissected tissue into an LC-MS-compatible buffer, leaving no need for solid-phase extraction or other methods for sample clean-up. For any contaminants associated with the FFPE tissue itself, we rely on the C18 material within an EvoTip Pure to act as a filter and use a disposable peptide trapping cartridge to ensure that material not compatible with LC-MS is not introduced to the analytical column.

We tested our workflow using different sample sizes of exocrine pancreas acinar tissue to assess its sensitivity and scalability. Additionally, we evaluated the protocol’s efficiency on cell populations isolated from tissue samples stained with conventional hematoxylin-eosin (H&E), which is commonly used in routine histopathological diagnostics, or selected based on their positivity for the epithelial cell adhesion molecule (EpCAM). This ensured that the protocol is adaptable to various staining methods and not dependent on any specific stain. With the established workflow, we then evaluated its applicability to, and retrievability of biological relevance from three types of cancer: pancreatic ductal adenocarcinoma (PDAC), lung adenocarcinoma (LAC), and glioblastoma (GB), where PDAC and LAC were Immunostained for EpCAM and GB was immunostained for the tumor-suppressor protein p53. We found distinct clustering between the three cancer types, along with prominent, cancer-type-specific protein expressions, demonstrating a proof-of-principle that our workflow is applicable for exploring biological questions.

## Materials and Methods

### Tissue Preparation

Resected tissue was fixed in 10% non-buffered formalin at a volume approximately 10 to 20 times that of the tissue for at least 24 hours and no longer than 72 hours, to mitigate over-crosslinking. After fixation, a pathologist macrodissected the surgical specimens, selecting tumor areas for paraffin embedding and further analysis. The tissue was dehydrated in successive concentrations of ethanol, increasing from 70% to 99%, cleared using Tissue-Clear®, and subsequently embedded in paraffin. Polyethylene naphthalate (PEN) membrane glass slides (Zeiss) were UV-treated for 1 hour, followed by overnight drying at 37 °C in a Binder drying and heating chamber. The tissue was then sectioned into 5 μm-thick slices using a rotary microtome and placed in a water bath at a temperature between 42 and 45 °C, depending on tissue composition, to “stretch” the paraffin sections and avoid folds and wrinkles. Following stretching, the tissue sections were collected onto the dried glass slide and left to dry at room temperature (RT).

### Hematoxylin-eosin (H&E) staining

After adhesion, the slides were deparaffinized by submerging them in Tissue-Tek® Tissue-Clear® Xylene Substitute for 10 minutes, followed by 20 dips in 99% ethanol, 10 dips in 96% ethanol, and finally, 10 dips in 70% ethanol. After deparaffinization, the ethanol was removed by submerging the slides in a vessel with running demineralized water for 5 minutes. Following the washing, the slides were submerged in Mayer’s hematoxylin for 4 minutes, running water for 5 minutes, 0.1% eosin in 0.1 M Walpole’s acetate buffer for 1 minute, and finally, running water for 3 minutes. The slides were then left in the fume hood to dry overnight, after which they were stored in a closed container at RT.

### EpCAM and p53 immunohistochemical staining

The day before staining, slides were heated overnight at 56 °C in a Binder drying oven, followed by 60 °C for 1 hour before staining. The tissue sections were deparaffinized in Tissue-Clear® for 10 minutes and rehydrated with decreasing concentrations of ethanol. The membrane slides for both EpCAM and p53 staining were transferred to automated immunostainers—Ventana Discovery for EpCAM and BenchMark Discovery Ultra (Ventana Medical Systems Inc.; Roche Diagnostics A/S, Hvidovre, Denmark) and Dako Omnis (Agilent Technologies) for p53. For EpCAM staining, antigen retrieval (pH 9.0) was performed in Ultra Cell Conditioning solution (CC1; Ventana) at 95 °C for 32 minutes, whereas in the p53 protocol, heat-induced epitope retrieval was conducted at 100 °C for 32 minutes using the same Tris-based CC1 buffer. Blocking of endogenous peroxidase in both protocols was achieved using Inhibitor CM (Ventana), though at different time points: in the EpCAM protocol, it was applied right before incubation with the primary antibody (Ab), while in the p53 protocol, it was performed after epitope retrieval for 8 minutes at RT. For EpCAM staining, sections were incubated with the EpCAM primary Ab (Ready-To-Use, Ventana) for 60 minutes at RT, followed by the Omnimap Anti-Mouse secondary Ab (Ventana). Detection was performed using the Discovery Chromomap DAB kit (Ventana). For p53 staining, sections were incubated with the p53 primary Ab (Bio-Rad Do7, MCA1703) at a 1:200 dilution, followed by DISCOVERY OmniMap anti-Ms HRP secondary Ab for 20 minutes at RT. Detection was completed using DISCOVERY DAB CM, DISCOVERY H2O2 CM, and DISCOVERY Copper CM. The slides were counterstained with Hematoxylin II for 8 minutes and Bluing Reagent for 4 minutes in both protocols—at RT for p53 and as part of the automated Ventana system for EpCAM.

### Sample collection by LCM

Twenty microliters of 100 mM triethylammonium bicarbonate (TEAB; Thermo Scientific™, Cat. No. 90114) in LiChrosolv® water (Supelco, Cat. No. 1153332500) were added to the lids of 0.5 mL Protein LoBind® tubes. We determined that 20 μL was the minimum volume capable of covering the entire surface of each lid, which is a critical factor when catapulting the tissue during LCM, as there is a high chance of losing samples if the tissue misses the buffer (however, it is possible to correct misaligned collections—where the tissue misses the droplet—by gently stirring the droplet with a pipette tip). The tubes were then individually loaded onto the ZEISS PALM Microbeam Laser Microdissection microscope using the SingleTube 500 CM II tube holder. During optimization, dense areas of exocrine pancreatic acinar cells between 1,166 μm^2^ and 800,000 μm^2^ with a thickness of 5 μm were collected in the buffer-filled lids, and collections were confirmed using a DinoLite digital microscope. For the validation of biological significance, ∼50,000 μm^2^ of PDAC, LAC, and GB cells were identified by trained histopathologists (C.V.R, E.S.R, B.W.K) and collected (PDAC and LAC samples included only type-specific cells. However, since GB tumor cells were dispersed as single cells within the stroma, these samples also contained stromal tissue.). Following the LCM collections, the tubes were spun down at max speed (for our centrifuge, this was 21,300 × g) for 5 minutes to ensure the tissue was released from the lids of the tubes.

### Sample processing

The samples were boiled at 95 °C for 5 minutes, followed by sonication for 10 cycles of 30 seconds on and 30 seconds off using a Bioruptor® Pico sonicator. The water was pre-cooled to 4.0 °C and maintained at this temperature throughout the sonication process (it is critical to ensure adequate cooling of the sonication bath, as higher temperatures can potentially damage the samples during sonication). After sonication, the samples were centrifuged at max speed for 30 seconds.

### Digestion

Ten μL of TEAB containing 1 ng of Trypsin/Lys-C Protease Mix (Pierce™ Thermo Scientific™, Cat. No. A41007) per 25,000 μm^2^ of tissue was added to the samples (to ensure optimal digestion kinetics, sample areas below 50,000 μm^2^ were given 2 ng of the digestion mix.). The samples were then digested overnight at 37 °C with rotational shaking at 1,000 rpm. The following day, the samples were centrifuged at max speed for 30 seconds. Note that we recommend adding 3 μL of 10% trifluoroacetic acid (TFA; Sigma-Aldrich, Cat. No. 302031-10×1ML) to the samples if they are to be stored for extended periods before analysis.

### EvoTip Loading

Activation of the C18 material within the EvoTips was performed by adding 20 *µ*L of LiChrosolv® acetonitrile (ACN; Supelco, Cat. No. 1000292500) to each tip and centrifuging at 700 x g for 1 minute (all centrifugation steps were conducted at 700 x g for 1 minute). The tips were then submerged in LiChrosolv® 2-Propanol (IPA; Supelco, Cat. No. 1027812500) for 1 minute. Following this, 20 *µ*L of LiChrosolv® water with 0.1% formic acid (Buffer A; Supelco, Cat. No. 1590132500) was added and centrifuged through. With the tips now activated, the samples were introduced and spun through. The tips were then spun through with 20 *µ*L of buffer A. Finally, 250 *µ*L of buffer A was added to the tips, and they were centrifuged for 10 seconds at 700 x g to ensure the liquid fully soaked the C18-bound samples. Additionally, approximately one-third of the EvoTip box was filled with buffer A to prevent sample drying.

### Liquid Chromatography settings

In our LC-MS/MS analysis, we employed the EvoSep One liquid chromatography system from Evosep Biosystems using the Whisper® 20 method. Peptides underwent separation over a 58-minute gradient (20 samples per day, SPD) at a flow rate of 100 nL/min. The column utilized was a 15 cm × 75 μm column with 1.7 μm C18 beads (Aurora Elite TS 15 cm nanoflow UHPLC column from IonOpticks). Mobile phases A and B comprised buffer A and LiChrosolv® acetonitrile with 0.1% formic acid (Buffer B; Supelco, Cat. No. 1590022500), respectively. The LC system was connected to an Orbitrap Eclipse Tribrid MS (Thermo Fisher Scientific) through an EasySpray ion source linked to a FAIMSPro device. An important notice: we experienced occasional, seemingly random LC overpressures, which we later discovered was caused by the accumulation of remaining FFPE debris on the sample collection needle. This was especially prominent for samples of smaller sizes, which appeared to aggregate more effectively. We found that this accumulation could be prevented by regularly (every 40–60 samples) wiping the sample needle, which often resulted in a visible collection of debris being removed.

### Mass spectrometry settings

For mass spectrometry data acquisition, we employed our in-house-developed WISH-DIA methods (16), which build on the HRMS1-DIA method by Xuan et al. 2020 (17). The FAIMSPro device was operated in positive mode, with a compensation voltage of -45 V. MS1 scans were performed at a resolution of 120,000, with automatic gain control (AGC) set to 300%, and the maximum injection time set to auto. The mass-to-charge ratio (m/z) range was specified from 400 to 1000. Higher energy collisional dissociation (HCD) was utilized for precursor fragmentation, with a normalized collision energy (NCE) of 33%, and the AGC target for MS2 was set to 1000%. Precursor ions were sequentially isolated in 17 m/z increments, scanned at a resolution of 120,000, with AGC set to 1000%, and the maximum injection time set to auto. Note that different methods, gradients (20 and 40 SPD), columns (Aurora and 15□cm × 75 μm ID column (PepSep)), and instruments (Orbitrap Eclipse and Orbitrap Exploris 480) were evaluated during the initial workflow development.

### Data analysis

The acquired raw MS data were analyzed using Spectronaut™ version 18 with the reviewed and filtered Homo sapiens library from UniProt, excluding isoforms. Spectronaut settings were mostly kept at factory configurations, except for “Enzymes / Cleavage Rules,” which were set to Trypsin and LysC. The fixed Carbamidomethyl (C) modification was removed, and “Quantity MS Level” was set to MS1. Searches and quantification were carried out in batches according to experimental conditions for all samples (binned by tissue amount, sample type, etc.), and follow-up data analysis was performed in the R environment (18,19).

## Results

### Developing a combined LCM and LC-MS/MS Workflow

For our initial approach, we optimized the protocol on exocrine pancreatic acinar cells and used commercially available LCM tubes with adhesive-filled caps. However, this method posed extensive challenges due to excessive tissue adherence, which prevented protein extraction. Therefore, inspired by Kwon et al. 2022 (10), we transitioned to a “hanging drop” method. This approach involves capturing the microdissected tissue in the lids of buffer-filled Protein LoBind tubes, effectively resolving the tissue-binding issues we encountered.

Next, we searched for an optimal collection and preparation buffer. We evaluated a range of compounds, including harsh lysis buffers such as guanidinium chloride (GdmCl) and hydrogen peroxide (H_2_O_2_), but observed improved yields from buffers not typically considered lysing agents, like PBS (phosphate-buffered saline) and TEAB. Additionally, we tested the effects of tris(2-carboxyethyl)phosphine hydrochloride (TCEP) and chloroacetamide (CAA), commonly used to reduce and alkylate disulfide bonds, respectively. Notably, the addition of TCEP and CAA for both TEAB and PBS lowered protein identification (TEAB+ and PBS+ in Figure 1). Based on these results, we proceeded with TEAB, which yielded slightly better outcomes than PBS. TEAB also has the advantage of being an LC-MS-compatible buffer.

**Figure 1.**
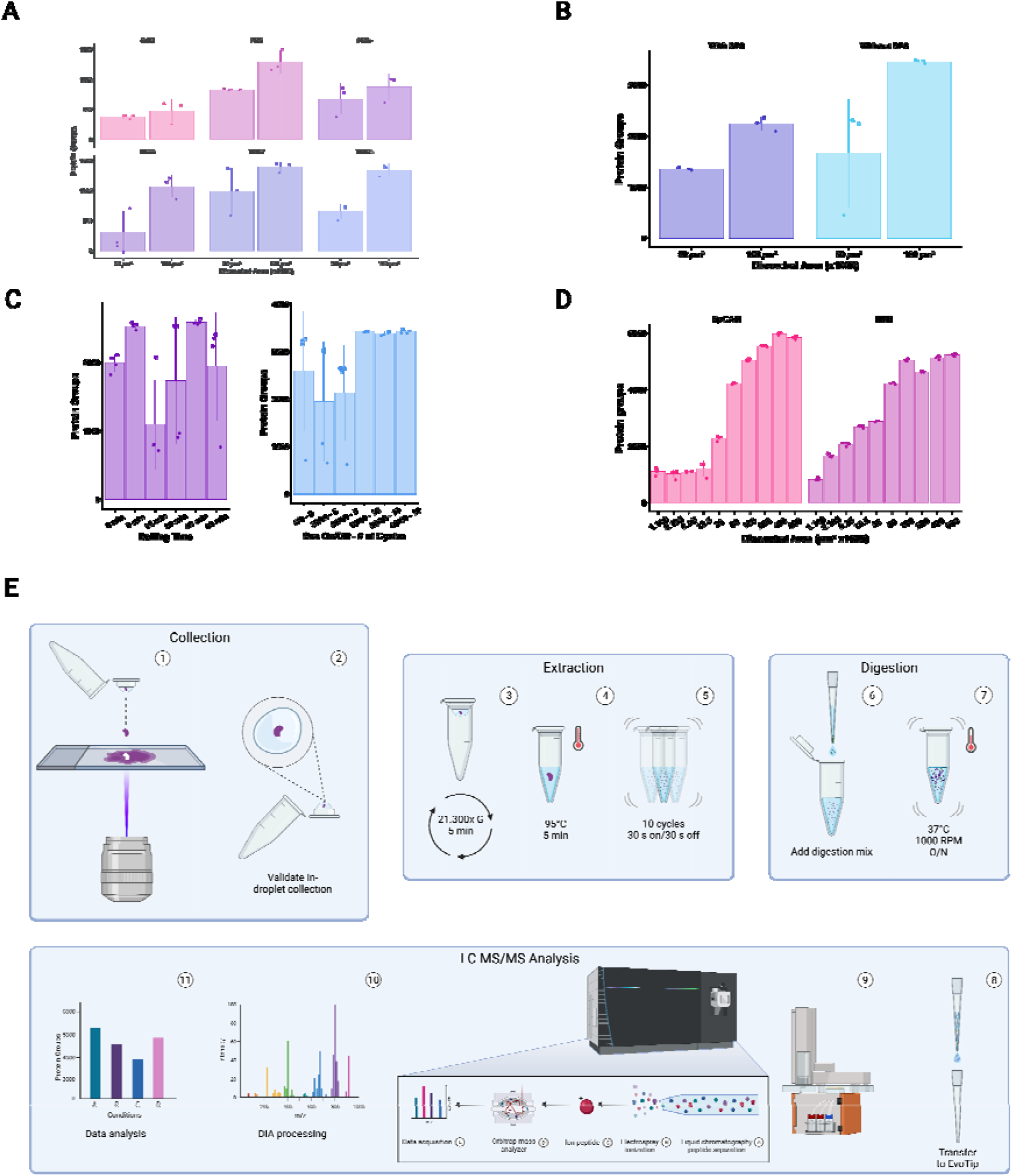
Workflow optimization overview on pancreatic acinar tissue. **(A)** Assessing different lysis buffers. TCEP and CAA were added to GdmCl, TEAB+, and PBS+. **(B)** Comparison between proteins yields with and without SP3 sample clean-up. **(C)** Evaluation of downstream sample processing of collected tissue (50,000 *µ*m^2^), with variations in boiling durations (on the left) and sonication periods and number of cycles (on the right). **(D)** Comparing our workflow on samples collected from tissue sections immunostained for EpCAM or stained with H&E, across a wide range of tissue sizes. Samples with less than 50% proteome coverage are removed. **(E)** Full overview of our optimized workflow for LC-MS/MS analysis of LCM-collected FFPE tissue.

From here, we assessed whether SP3 clean-up could be omitted, as TEAB introduces no contaminating elements (20). Omitting SP3 clean-up resulted in an increase of ∼55% at 50,000 *µ*m^2^ and ∼70% at 100,000 *µ*m^2^ of material in identified proteins, achieving similar depths with only half the amount of tissue (Figure 1). Importantly, no negative impact on column performance was observed, suggesting that we can successfully mitigate sample losses associated with SP3 clean-up. Subsequently, we evaluated downstream processes, particularly the duration of tissue boiling and the impact of sonication (Figure 1). Testing different boiling durations revealed that periods beyond 5 minutes had no effect on the number of identified proteins, while the difference between no boiling and 5 minutes resulted in a ∼25% increase. Sonication showed little to no effect on protein yields; however, reproducibility increased significantly for samples sonicated at 10 cycles. Furthermore, we examined the impact of trypsin concentration, the volume of the collection droplet, and protein depth when using the Thermo Orbitrap Astral Mass Spectrometer instead of an Eclipse Tribrid—results are detailed in the Supplementary Data.

With our optimized workflow, we evaluated performance across varying tissue amounts and staining methods, assessing proteomic depth while preserving spatial localization. EpCAM-stained tissue showed slightly higher performance compared to tissue dissected from H&E-stained sections, but we cannot exclude that these results originate from the fact that different tissue sections were used. Both staining methods approached a plateau around ∼200,000 μm^2^, suggesting an optimal balance between low input and high protein identification at ∼50,000 μm^2^, or an estimated 82 cells by volume (calculated by assuming the cells are spherical and are similar in size to acinar rat cells, which have a reported average diameter of 18 *µ*m (21)) (Figure 1).

### Evaluating biological significance

As a proof of concept, we aimed to evaluate the applicability of our workflow across three different cancer types and assess its biological relevance. We isolated groups of cancer cells from three malignancies: Pancreatic Ductal Adenocarcinoma (PDAC), Glioblastoma (GB), and Lung Adenocarcinoma (LAC). Representative amounts of cells from each cancer type (50,000 *µ*m^2^) were isolated and subjected to our sample preparation and LC-MS acquisition.

We observed similar numbers of identified proteins across the different tissue types, demonstrating that the workflow is versatile and applicable to a wide range of tissues (Figure 2, Figure ***2***). Both the PCA plot and the hierarchical heatmap revealed distinct tissue clustering (Figure 2, Figure ***2***), underscoring the ability of the workflow to capture biological relevance. When overlaid with the volcano plot (Figure 2), distinct protein signatures known to be upregulated in the respective tissues were evident, such as TPPP3, which is expressed in glial cells and the respiratory system, and TUBB4A, MAP1B, and FMNL2, which are abundantly expressed in the brain (according to The Human Protein Atlas, (22)). Demonstrating the workflow’s ability to identify biological differences between distinct tissue types.

**Figure 2.**
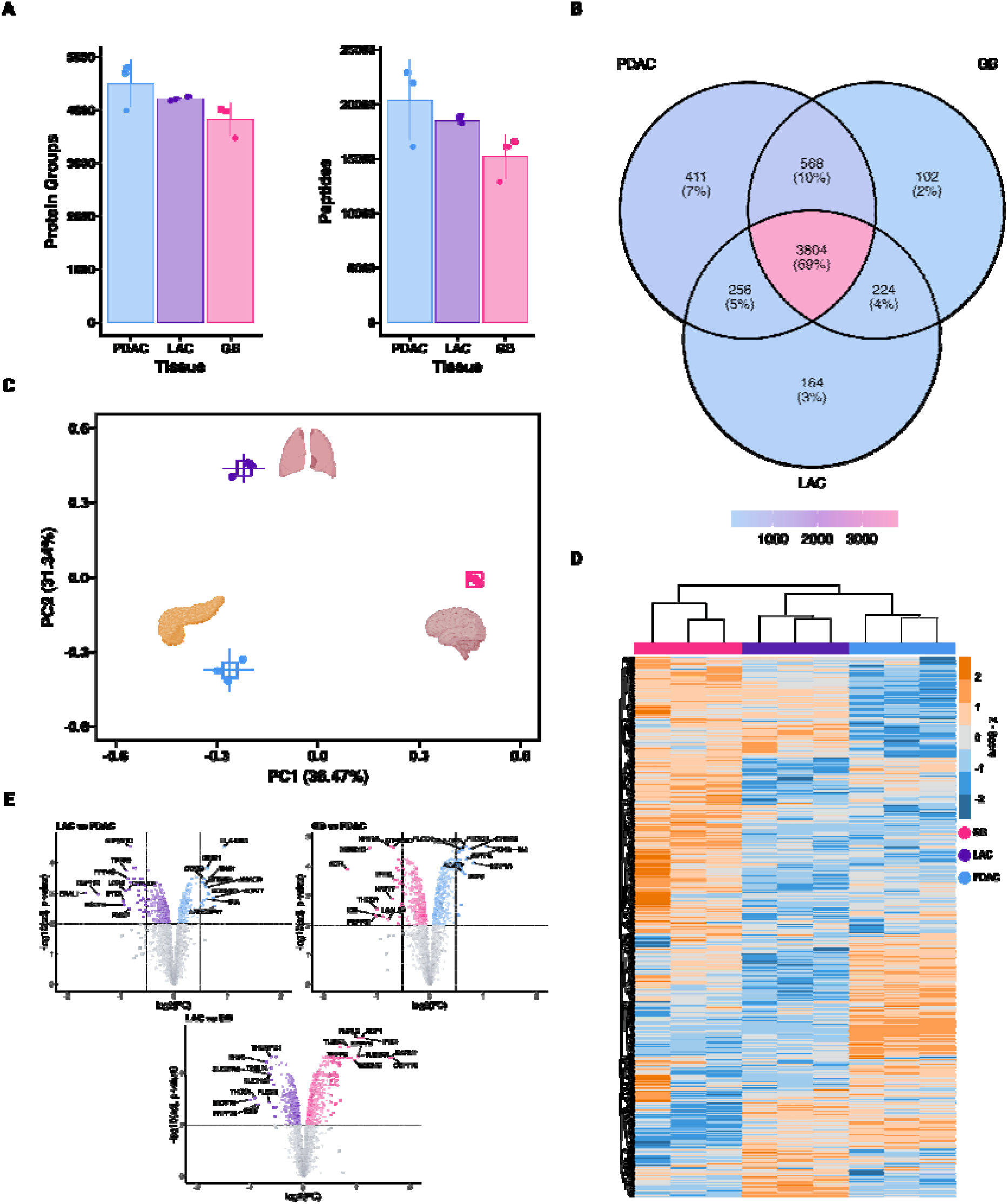
Assessing biological insights across various cancer types. **(A)** Average number of protein groups (left) and peptides (right) detected from the three cancer types (PDAC, LAC, and GB). **(B)** Venn diagram showing the number of commonly and uniquely expressed proteins, in the three different cancer types. **(C)** PCA plot showing clustering of PDAC (blue), LAC (purple), and GB (pink). **(D)** Heatmap showing relative up-regulation (orange) and down-regulation (blue) of protein levels between the three cancer types. **(E)** Volcano plots comparing differential expression in the three cancer types, with the top 10 mostly differentially expressed proteins labelled.

Lastly, we evaluated the potential for drug-target and biomarker discoveries using our workflow. Since most drugs target epigenetic regulators, kinases, transcription factors, or surface proteins, we compared the identified proteins across the different cancers against such libraries, finding a solid array of hits. Figure 3 and Figure ***3***, illustrate the number of unique protein targets identified in each cancer type, highlighting their potential relevance. Among these, we focused specifically on the kinases uniquely detected in PDAC and subjected them to Gene Ontology (GO) enrichment analysis (Figure 3). The analysis revealed a diverse array of enriched pathways, including positive regulation of macrophage proliferation, beta-catenin destruction complex, protein kinase complex assembly, and type B pancreatic cell development, with the latter again showing that we can identify tissue-specific signatures (23).

**Figure 3.**
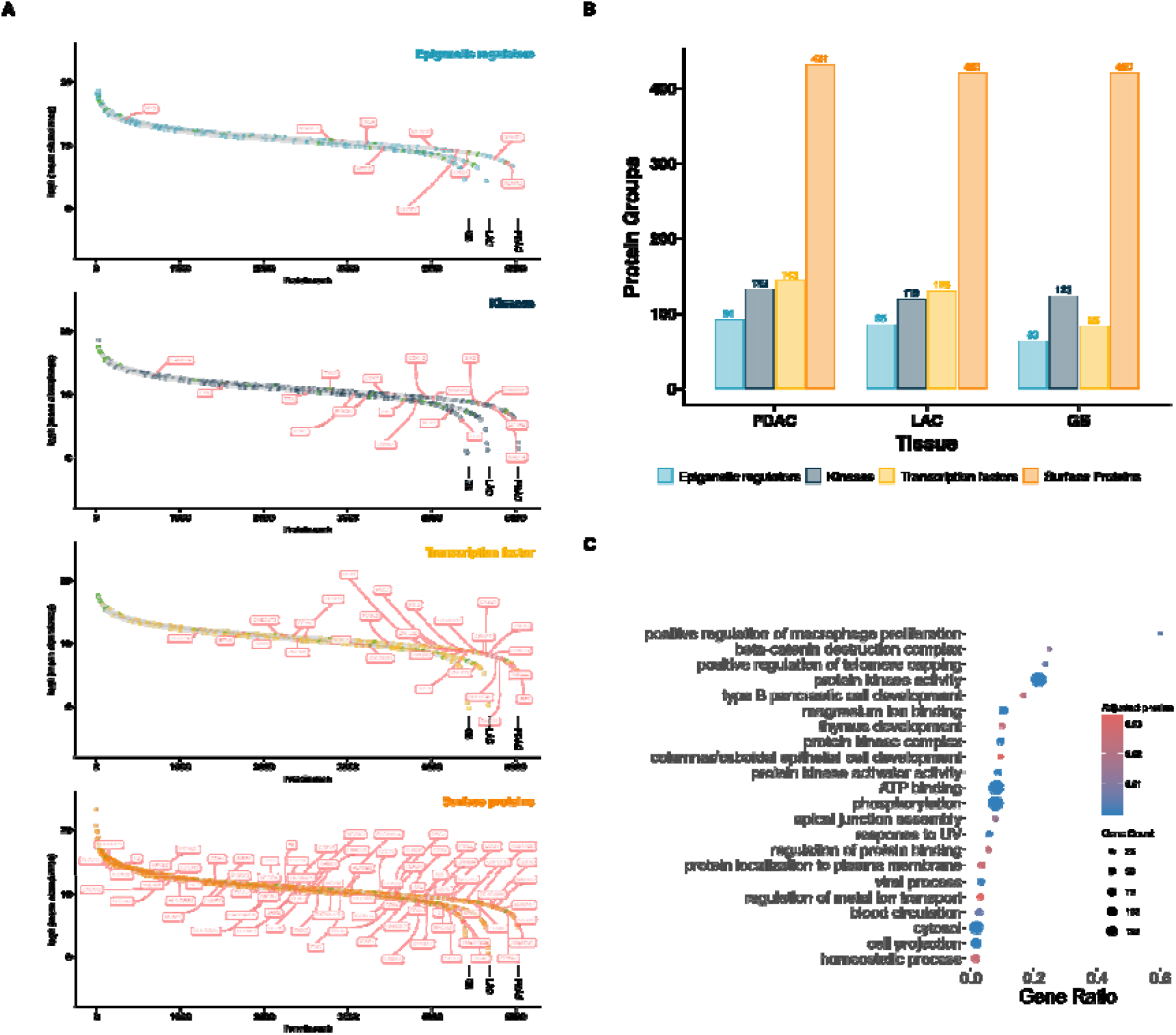
Evaluating our workflow for drug-target and biomarker discovery. **(A)** Ranked plots of the three cancer types (PDAC, LAC, and GB) with highlighted protein categories. Proteins unique to each tissue type are highlighted and labeled in red, while known biomarkers related to each specific tissue type are highlighted in green. **(B)** Total number of proteins detected within the four protein categories for each cancer type. **(C)** GO enrichment performed on kinases identified only within PDAC samples.

## Discussion

In this study, we have developed an LC-MS-compatible LCM-based sample preparation workflow that facilitates deep proteome coverage from archival tissue, without the need for libraries such as high-pH or high-load. While multiple workflows for the Leica LMD series exist, the more modest ZEISS PALM system has lacked robust protocols. By adapting the hanging drop method, we provide a straightforward and effective solution for spatial proteomics, ensuring reliable tissue collection and maximizing protein identification. Our approach builds upon and extends previous efforts in spatial proteomics, demonstrating comparable or improved performance relative to established methods from leading laboratories, such as those of Matthias Mann (11), Fabian Coscia (13), and Ryan Kelly and Ying Zhu (10,24). Unlike conventional workflows that require high material input, our method enables deep proteome coverage from as few as 80 cells, making it an accessible and reproducible approach for FFPE tissue analysis. Furthermore, we observed that staining with either standard H&E or EpCAM immunostaining for optimal detection of epithelial tumor cells had no significant impact on protein retrieval, reinforcing the robustness of our workflow across different sample preparation conditions.

A major advantage of our method is its simplicity and cost-effectiveness, eliminating unnecessary processing steps while improving protein identification. Additionally, our external quality control step using a DinoLite digital microscope, enhanced sample reproducibility by allowing us to detect and correct inefficiencies in tissue collection. However, challenges remain, particularly in achieving high-throughput capabilities, where the implementation of instruments such as Cellenone, as demonstrated by Mahmut et al., 2024 (14), could greatly improve throughput. Additionally, our study was conducted on a limited number of patient samples, and further validation across a broader cohort will be necessary to fully establish the method’s generalizability.

Future applications of this workflow may enable even finer spatial resolution, extending its use to small tissue biopsies in routine histopathological diagnostics and advancing spatial single-cell proteomics by MS (scp-MS), as shown in recent studies such as the one by Rosenberger et al., 2023 (25). By continuing to refine sample preparation strategies and integrating with emerging analytical techniques, this approach has the potential to unlock new insights into tissue heterogeneity, cancer biology, and biomarker discovery.

## Conclusion

In conclusion, our workflow for archival tissue represents a simple approach that effectively addresses the challenges associated with FFPE samples and tissue heterogeneity. Designed for adaptability and efficiency, and relying solely on off-the-shelf consumables, this method unlocks the potential of low-input materials for retrospective proteomics studies, making extensive use of FFPE repositories. While most of the best efforts presented by others so far have primarily been conducted on the timsTOF platform from Bruker, we instead evaluated the capability of Orbitrap-based instruments to achieve similar low-input, spatial proteomics workflows. Across diverse cancer types, this workflow demonstrates its ability to unravel intricate proteome landscapes and molecular nuances within each microenvironment, making it well-suited for data-driven investigations of cell-type-specific proteome dynamics in primary clinical specimens.

## List of abbreviations

Ab: Antibody
AGC: Automatic Gain Control
CAA: Chloroacetamide
DIA: Data Independent Acquisition
EpCAM: Epithelial Cell Adhesion Molecule
FFPE: Formalin-fixed, paraffin-embedded
GB: Glioblastoma
GdmCl: Guanidinium Chloride
GO: Gene Ontology
H&E: Hematoxylin and Eosin
HCD: Higher energy collisional dissociation
IPA: Isopropanol / 2-Propanol
LAC: Lung Adenocarcinoma
LC-MS/MS: Liquid Chromatography-Mass Spectrometry/Mass Spectrometry
LCM: Laser Capture Microdissection
NCE: Normalized collision energy
PBS: Phosphate Buffered Saline
PDAC: Pancreatic Ductal Adenocarcinoma
RT: Room Temperature
SPD: Samples Per Day
SP3: Single-Pot, Solid-Phase-enhanced Sample Preparation
TCEP: Tris(2-carboxyethyl)phosphine hydrochloride
TEAB: Triethylammonium Bicarbonate
TFA: Trifluoroacetic Acid
WISH-DIA: Wide Isolation window High-resolution MS1-DIA

## Data availability

The datasets supporting the conclusions of this article are available in the PRIDE Proteomics Identification Database.

https://www.ebi.ac.uk/pride/archive/projects/PXD060512

## Declarations

### Ethics approval

Surgical specimens of human pancreatic exocrine acinar tissue and adenocarcinoma (PDAC), lung adenocarcinoma (LAC), and glioblastoma (GB) were obtained from routine resections performed at Rigshospitalet, Copenhagen University Hospital, after receiving approval from the Ethics Committee prior to the initiation of the project (Pancreas: J. no: VD-2018-446, I-suite no. 6701; Brain: VEK: S-20150148; Lung: VEK: H-23021044). Additionally, the Danish Registry for Use of Tissue was checked to ensure that patients included in the study had not refused the usage of their tissue samples for research purposes.

### Consent for publication

Not applicable.

### Author contributions

R.D., E.M.S., and P.K. conceived and designed the project. R.D., P.K and E.M.S. performed the experimental investigations. C.V.R., P.R.K., E.S.R., and B.W.K selected tumor areas in the FFPE specimens. N.S.B, K.S, and P.R.K aided in sectioning and staining of FFPE samples. Data analysis was performed by R.D, J.W, and V.P. The original manuscript was drafted by R.D., with all other authors contributing to review and editing. E.M.S. and P.K supervised the work.

## Acknowledgements

We would like to thank all members from the Cell Diversity Lab, headed by E.M.S. for constructive input and fruitful discussions, and the DTU Proteomics Core for technical instrument support.

## References

1. Kokkat, T. J., Patel, M. S., McGarvey, D., LiVolsi, V. A., & Baloch, Z. W. (2013). Archived Formalin-Fixed Paraffin-Embedded (FFPE) Blocks: A Valuable Underexploited Resource for Extraction of DNA, RNA, and Protein. Biopreservation and Biobanking, 11(2), 101–106. 10.1089/bio.2012.0052

2. Eccher, A., Scarpa, A., & Dei Tos, A. P. (2023). Impact of a centralized archive for pathology laboratories on the health system. Pathology Research and Practice, 245, 154488. 10.1016/j.prp.2023.154488

3. Wiśniewski, J. R., Duś, K., & Mann, M. (2013). Proteomic workflow for analysis of archival formalin-fixed and paraffin-embedded clinical samples to a depth of 10000 proteins. Proteomics - Clinical Applications, 7(3-4), 225–233. 10.1002/prca.201200046

4. Ramón y Cajal, S., Sesé, M., Capdevila, C., Aasen, T., De Mattos-Arruda, L., Diaz-Cano, S. J., … Castellví, J. (2020). Clinical implications of intratumor heterogeneity: challenges and opportunities. Journal of Molecular Medicine, 98(2), 161–177. 10.1007/s00109-020-01874-2

5. Specht, H., & Slavov, N. (2018). Transformative Opportunities for Single-Cell Proteomics. Journal of Proteome Research, 17(8), 2565–2571. 10.1021/acs.jproteome.8b00257

6. Janku, F. (2014). Tumor heterogeneity in the clinic: Is it a real problem? Therapeutic Advances in Medical Oncology, 6(2), 43–51. 10.1177/1758834013517414

7. Zhu, L., Jiang, M., Wang, H., Sun, H., Zhu, J., Zhao, W., … He, Y. (2021). A narrative review of tumor heterogeneity and challenges to tumor drug therapy. Annals of Translational Medicine, 9(16), 1351. 10.21037/atm-21-1948

8. Valdemaras Petrosius, Pedro Aragon-Fernandez, Tabiwang N. Arrey, Nil Üresin, Benjamin Furtwängler, Hamish Stewart, Eduard Denisov, Johannes Petzoldt, Amelia C. Peterson, Christian Hock, Eugen Damoc, Alexander Makarov, Vlad Zabrouskov, Bo T. Porse, Erwin M. Schoof: Evaluating the capabilities of the Astral mass analyzer for single-cell proteomics. bioRxiv. Preprint. 10.1101/2023.06.06.543943

9. Brunner, A. D., Thielert, M., Vasilopoulou, C., Ammar, C., Coscia, F., Mund, A., … Mann, M. (2022). Ultra-high sensitivity mass spectrometry quantifies single-cell proteome changes upon perturbation. Molecular Systems Biology, 18(3), e10798. 10.15252/msb.202110798

10. Kwon, Y., Piehowski, P. D., Zhao, R., Sontag, R. L., Moore, R. J., Burnum-Johnson, K. E., … Zhu, Y. (2022). Hanging drop sample preparation improves sensitivity of spatial proteomics. Lab on a Chip, 22(15), 2869–2877. 10.1039/d2lc00384h

11. Mund, A., Coscia, F., Kriston, A., Hollandi, R., Kovács, F., Brunner, A. D., … Mann, M. (2022). Deep Visual Proteomics defines single-cell identity and heterogeneity. Nature Biotechnology, 40(8), 1231–1240. 10.1038/s41587-022-01302-5

12. Davis, S., Scott, C., Oetjen, J., Charles, P. D., Kessler, B. M., Ansorge, O., & Fischer, R. (2023). Deep topographic proteomics of a human brain tumour. Nature Communications, 14(1), 7710. 10.1038/s41467-023-43520-8

13. Makhmut, A., Qin, D., Fritzsche, S., Nimo, J., König, J., & Coscia, F. (2023). A framework for ultra-low-input spatial tissue proteomics. Cell Systems, 14(11), 1002–1014.e5. 10.1016/j.cels.2023.10.003

14. Anuar Makhmut, D. Qin, David Hartlmayr, Anjali Seth, Fabian Coscia: An automated and fast sample preparation workflow for laser microdissection guided ultrasensitive proteomics. bioRxiv. Preprint. 10.1101/2023.11.29.569257

15. Chen, H., Zhang, Y., Zhou, H., Chen, W., Peng, J., Feng, Y., … Liu, S. (2023). Routine Workflow of Spatial Proteomics on Micro-formalin-Fixed Paraffin-Embedded Tissues. Analytical Chemistry, 95(45), 16733–16743. 10.1021/acs.analchem.3c03848

16. Petrosius, V., Aragon-Fernandez, P., Üresin, N., Kovacs, G., Phlairaharn, T., Furtwängler, B., … Schoof, E. M. (2023). Exploration of cell state heterogeneity using single-cell proteomics through sensitivity-tailored data-independent acquisition. Nature Communications, 14(1), 5910. 10.1038/s41467-023-41602-1

17. Xuan, Y., Bateman, N. W., Gallien, S., Goetze, S., Zhou, Y., Navarro, P., … Conrads, T. P. (2020). Standardization and harmonization of distributed multi-center proteotype analysis supporting precision medicine studies. Nature Communications, 11(1), 5248. 10.1038/s41467-020-18904-9

18. Ritchie, M. E., Phipson, B., Wu, D., Hu, Y., Law, C. W., Shi, W., & Smyth, G. K. (2015). Limma powers differential expression analyses for RNA-sequencing and microarray studies. Nucleic Acids Research, 43(7), e47. 10.1093/nar/gkv007

19. Szklarczyk, D., Kirsch, R., Koutrouli, M., Nastou, K., Mehryary, F., Hachilif, R., … Von Mering, C. (2023). The STRING database in 2023: protein-protein association networks and functional enrichment analyses for any sequenced genome of interest. Nucleic Acids Research, 51(1 D), D638–D646. 10.1093/nar/gkac1000

20. Hughes, C. S., Moggridge, S., Müller, T., Sorensen, P. H., Morin, G. B., & Krijgsveld, J. (2019). Single-pot, solid-phase-enhanced sample preparation for proteomics experiments. Nature Protocols, 14(1), 68–85. 10.1038/s41596-018-0082-x

21. Morgan, R. G., Schaeffer, B. K., & Longnecker, D. S. (1986). Size and number of nuclei differ in normal and neoplastic acinar cells from rat pancreas. Pancreas, 1(1), 37–43. 10.1097/00006676-198601000-00008

22. https://www.proteinatlas.org/

23. Liis Kolberg, Uku Raudvere, Ivan Kuzmin, Priit Adler, Jaak Vilo, Hedi Peterson: g:Profiler—interoperable web service for functional enrichment analysis and gene identifier mapping (2023 update) Nucleic Acids Research, May 2023; doi:10.1093/nar/gkad347

24. Woo, J., Williams, S. M., Markillie, L. M., Feng, S., Tsai, C. F., Aguilera-Vazquez, V., Sontag, R. L., Moore, R. J., Hu, D., Mehta, H. S., Cantlon-Bruce, J., Liu, T., Adkins, J. N., Smith, R. D., Clair, G. C., Pasa-Tolic, L., & Zhu, Y. (2021). High-throughput and high-efficiency sample preparation for single-cell proteomics using a nested nanowell chip. Nature Communications, 12(1), 6246. 10.1038/s41467-021-26514-2

25. Rosenberger, F. A., Thielert, M., & Mann, M. (2023). Making single-cell proteomics biologically relevant. Nature Methods, 20(3), 320–323. 10.1038/s41592-023-01771-9

26. Illustrations were created with BioRender.com

