## Supplementary Figure 1 and 2 for "Refining Spatial Proteomics by Mass Spectrometry: An Efficient Workflow Tailored for Archival Tissue"

Supplemntary

During method development, we also tested the impact of different collection volumes and trypsin concentrations on protein yield and identification rates. To optimize digestion efficiency, we compared trypsin-to-protein ratios ranging from 1:50 to 10:1 and observed that concentrations above 1:2.5 did not significantly improve protein identifications. Similarly, we evaluated collection volumes between 5 µL and 40 µL per laser capture site, finding that volumes below 20 µL result in lower numbers and, more importantly, lower reproducibility (Figure 1).

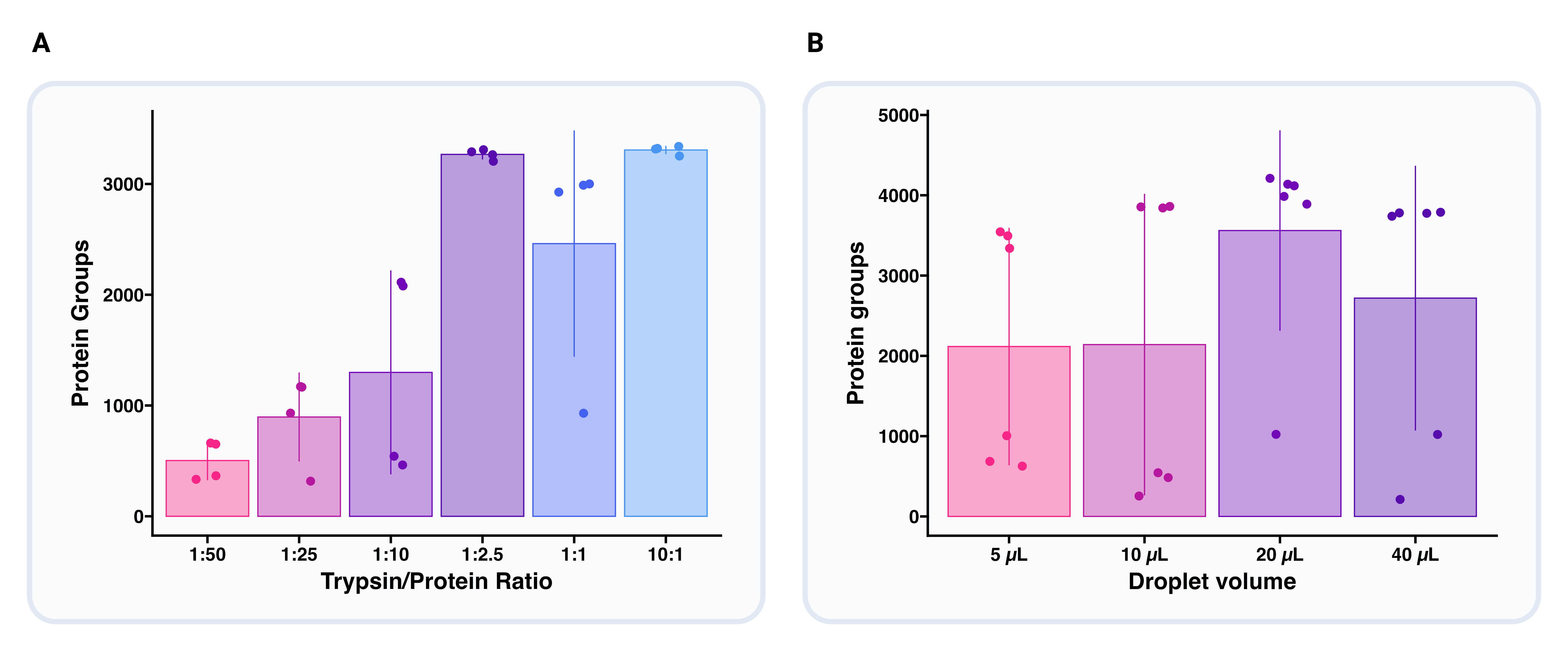

**Figure 1. (A)** Yield comparison between different volumes of LCM collection droplets. **(B)** Yield comparison between different trypsin concentrations.

Lastly, we compared the capabilities of the Orbitrap Astral and the Orbitrap Eclipse mass spectrometers. The Astral improves yields from 50,000 µm² of pancreatic acinar cells by approximately 64% at 20 SPD and 69% at 40 SPD, compared to the Eclipse (Figure 2).

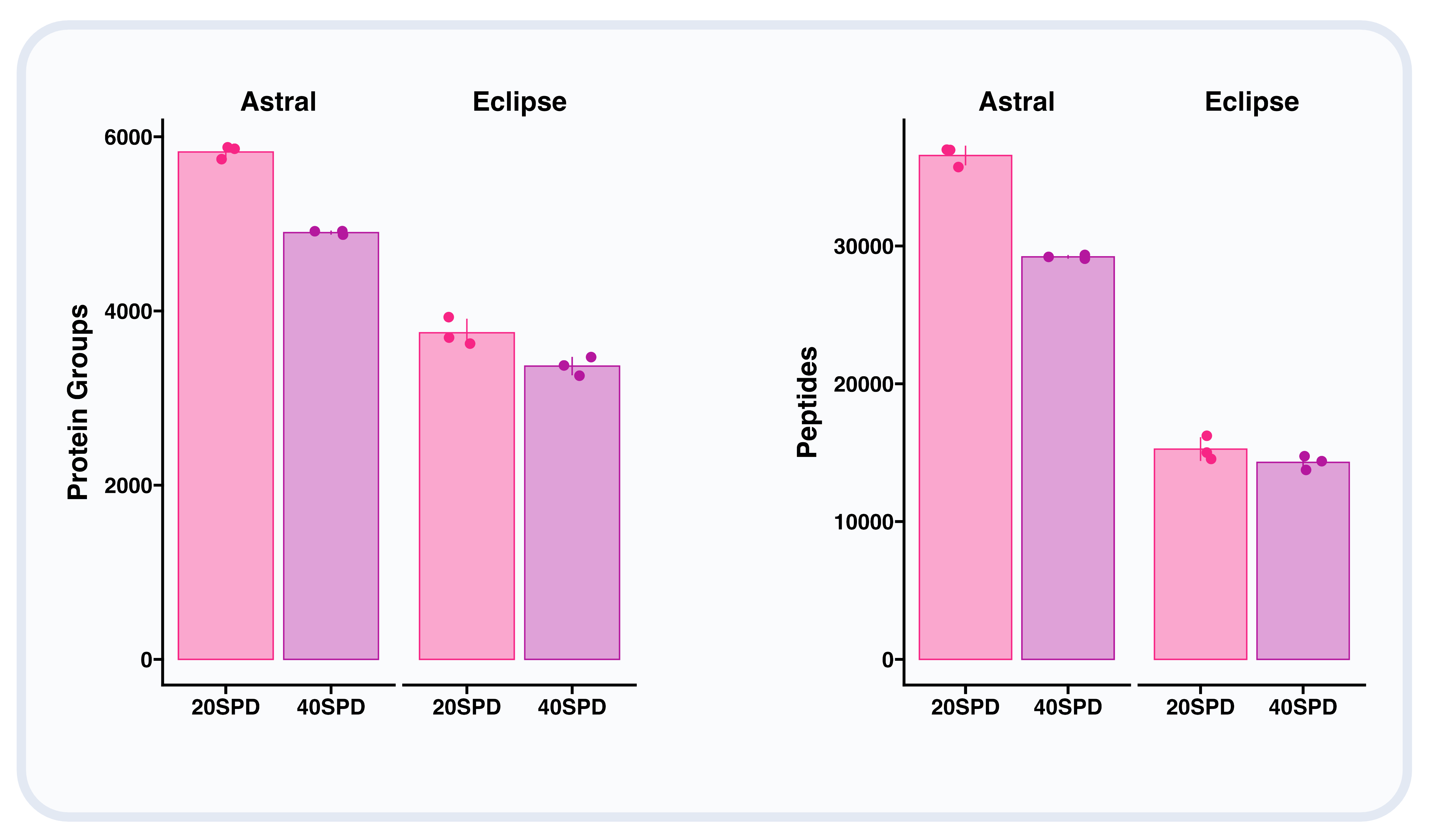

**Figure 2.** Number of protein groups and peptides detected using the Orbitrap Astral and Orbitrap Eclipse at 20- and 40-SPD gradients.
